# Elevated Activity in Left Homologous Music Circuits is Maladaptive for Music Perception but Mediated by Decoupled Structure and Function

**DOI:** 10.1101/2024.02.04.578219

**Authors:** Yucheng Wang, Zhishuai Jin, Sizhu Huyang, Qiaoping Lian, Daxing Wu

**Affiliations:** Medical Psychological Center, The Second Xiangya Hospital, Central South University, Changsha, Hunan, 410011, China; Medical Psychological Institute of Central South University, Changsha, Hunan, 410011, China; National Clinical Research Center on Mental Disorders, Changsha, Hunan, 410011, China

**Keywords:** structure-function coupling, amusia, music perception, regional activity, contralateral homologous region

## Abstract

Music is inherent in human life and is a significant topic of cognitive neuroscience. Previous studies focused on amusia suggested that two frontotemporal circuits engage in music processing. Structure-function coupling is an important feature of human brain, which is associated with cognition and allows for a more sensitive investigation of brain-behavior association. However, we still have limited knowledge about the relation between structure-function coupling, music processing and other regional neural profiles. We recruited 106 participants (43 subjects were diagnosed with congenital amusia) and measured their music perception by Montreal Battery of Evaluation of Amusia (MBEA). Then we utilized support vector regression algorithm and mediation analysis, and employed amplitude of low frequency fluctuation (ALFF), structural/functional degree centrality (DC) and structure-function coupling to explore their relation with global averaged MBEA score. We found structure-function coupling of widespread brain regions in both hemispheres, rather than ALFF or DC, contributed to predict MBEA score. Left middle frontal gyrus, bilateral inferior temporal gyrus and right insula were most predictive regions, and these regions were involved in memory and cognitive control according to meta-analysis. Further, coupling of left middle frontal gyrus, a region that is homologous to and is connected with typical music circuits, fully mediated the negative relation between ALFF and MBEA score. Our findings provide further understanding for the neural basis of music, and have implications for neural plasticity, neuromodulation therapy and cognitive causes of amusia.

**Highlights:**

- Our study first applies structure-function coupling to investigate the neural correlates of music perception, and predicting modeling indicate structure-function coupling is more effective than regional activity and connectivity.
- Elevated activity of contralateral homologous of music circuits might be maladaptive rather than compensatory.
- Elevated spontaneous regional activity disrupts their connections, which might be a specific expression of neural plasticity for certain regions.
- Our findings have implications for cognitive causes (abnormal memory and/or cognitive control to music salients) of amusia.

## 1. Introduction

Music is a fundamental but important element in human life. It serves as a means of social communication and contributes to mental health during stressful periods (Mas-Herrero et al., 2023). However, there is a specific neurological impairment called amusia, which is characterized by music perception and/or production deficits but could not be explained by general cognitive or motor deficits. Neuroimage studies focused on amusia contribute to our understanding of neural mechanisms of music and published articles have provided some insights. Generally, two neural frontotemporal feedback circuits: right arcuate fasciculus (AF)-based dorsal stream and right inferior fronto-occipital fasciculus (IFOF)-based ventral stream engaged in music cognition (Peretz, 2016; Sihvonen et al., 2019). In which, right superior temporal gyrus (STG) and inferior frontal gyrus (IFG) are core regions, other regions such as the right insula and inferior parietal lobule (IPL) located on both circuits also play a role in music processing. Altered function or structure in both circuits were reported (Albouy et al., 2019; Albouy et al., 2013; Chen and Yuan, 2016; Hyde et al., 2007; Jin et al., 2021a; Liao et al., 2022; Norman-Haignere et al., 2016; Sihvonen et al., 2016; Sihvonen et al., 2017a; Sihvonen et al., 2021; Sihvonen et al., 2017b; Sun et al., 2021). There is an acknowledged cognition-related interaction between structure and function, however, current studies only explored musical neural processes via multiple unimodal features separately to date. The association how connect with other neural profiles and affect music processing needs further discussion.

Structure-function coupling, a measure of structure-function association, describes structural support for function communication. Previous studies indicated structure-function coupling generally decreases both strength and heritability following the hierarchical distribution from primary sensory-motor cortex to higher order multimodal cortex (Baum et al., 2019; Gu et al., 2021; Liu et al., 2023b; Valk et al., 2022; Vázquez-Rodríguez et al., 2019), and is related to multiple cognitive functions or clinical outcomes (Baum et al., 2019; Cao et al., 2020; Chan et al., 2022; Chen et al., 2022; Collin et al., 2017; Koubiyr et al., 2019; Kulik et al., 2022; Liu et al., 2023a; Pan et al., 2023; Tay et al., 2023; Wang et al., 2018; Wu et al., 2023; Zarkali et al., 2021; Zhang et al., 2019; Zhang et al., 2022). Moreover, it is argued that connectivity patterns of certain brain areas or circuits are central to understanding their function whereas structure-function coupling combines the principles of functional organization (segregation and integration) and both connectivity information (Genon et al., 2018; Tononi et al., 1994), which suggested structure-function coupling may allow for a more effective detection of brain-behavior association. Indeed, some research support the opinion (Litwińczuk et al., 2022; Zhang et al., 2011). Therefore, we argued that structure-function coupling is more effective than other regional neural profiles when capturing the neural correlates of music.

In current study, we aim to investigate the relation between structure-function coupling and music perception, and whether structure-function coupling could better capture neural correlates of music than regional activity and connectivity. We hypothesized that: 1) music perception difference among adults is associated with variant structure-function coupling; 2) structure-function coupling is a more effective neural feature when explaining music perception. Additionally, we would conduct exploratory analysis to examine the multi-relationship between music perception and structure-function coupling, regional activity and connectivity.

## 2. Methods

### 2.1. Participants

All participants were recruited from Chinese universities in Hunan via campus posters from 2018 to 2022. Inclusion criteria include: 1) right-handed; 2) music training-naïve. Exclusion criteria include: 1) abnormal hearing measured by pure tone audiometry; 2) history of mental or neurological disorders; 3) Wechsler Adult Intelligence Scale (WAIS; IQ) score lower than 85; 4) contraindications for magnetic resonance imaging (MRI). Finally, 107 participants were recruited and all signed inform consent. The Ethics Committee of the Second Xiangya Hospital, Central South University approved the research.

### 2.2. Music Perception Measures

The Montreal Battery of Evaluation of Amusia (MBEA) is an assessment to recognition CA (Peretz et al., 2003). MBEA includes 6 subtests (scale, contour, interval, rhythm, meter and memory), and each subtest consists of 30 valid items. Participants need to determine whether same or not of paired music, the type of specific music, or whether appeared in first 5 subtests. Global averaged score of all subtests was calculated and used as quantitative evaluation of music perception. According to previous study (Nan et al., 2010), individuals with MBEA score lower than 21.5 were considered as CA.

### 2.3. MRI Data Acquisition

All MRI data were collected using a 3T Siemens Skyra scanner at the Second Xiangya Hospital, Central South University. Resting-state functional MRI images and T1-weighted structural images were obtained by echo-planar imaging and magnetization-prepared rapid gradient echo sequence, detailed scanning parameters were reported in our published articles (Jin et al., 2021a; Jin et al., 2021b). Diffusion MRI was acquired along 64 directions (b = 1000s/mm^2^) together with non-diffusion weighting (b = 0s/mm^2^) with following parameters: repetition time = 7000 ms, echo time = 86 ms, slices = 60, slice thickness = 2 mm, acquisition matrix = 128 × 128, flip angle = 90° and voxel size = 2 × 2 × 2.5 mm^3^.

### 2.4. MRI Data Preprocessing

DPABI 7.0 (Yan et al., 2016) and FSL 6.0.6 was used to preprocess functional and diffusion MRI data separately. For functional data, main preprocessing steps included: removing first 10 time points, slice timing correction, realign, spatial normalization (resampled to 3 × 3 × 3 mm), smoothing (6mm smoothing kernel), nuisance regression (linear trend, Friston-24 head motion parameters, cerebrospinal fluid and whiter matter signals) and band pass filter (0.01 – 0.1 Hz). For diffusion data, main processing steps included eddy-current and head motion correction, gradient direction correction, bet and get brain mask from b0 image, calculating fractional anisotropy (FA). One participant was removed due to mean frame-wise displacement (FD) > 0.2, 106 participants were included in subsequent analysis.

### 2.5. Functional and structural connectome

For each subject, functional and structural connectivity matrices were defined using the Brainnetome Atlas (Fan et al., 2016) which parcellated cerebral cortex into 246 regions of interests (ROI). Person correlation of time series between each pair of ROIs was calculated and generated 246 × 246 symmetric matrices. Fisher’s Z transformation was applied to the matrix representing functional connectivity. *BedpostX* and *probtrackX2* commands in FSL with default parameters (5000 samples were initiated in each seed voxel) were used to conduct probabilistic tractography. Original asymmetric matrices were transformed to symmetric form with following formula: (S_ij_ + S_ji_) / 5000 × (N_i_ + N_j_), S_ij_ is original streamline counts that streamlines seeded in ROI_i_ and reached ROI_j_, N_i_ is volume of ROI_i_. Further, a consistency-based method (coefficient of variation of each edge) was applied (Roberts et al., 2017) to reduce spurious connections induced by probabilistic tractography because the criteria contribute to retain consistent both short- and long-range connections across subject. According to previous studies (Buchanan et al., 2020; Roberts et al., 2017), a 30^th^ percentile was used to remove inconsistent (spurious) structural probabilistic connections.

### 2.6. Structure-function Coupling, ALFF and Degree Centrality

Spearman correlation coefficient between non-zero structure connectivity and corresponding functional connectivity represent structure-function coupling of each ROI. Amplitude of low frequency fluctuation (ALFF), a measure of regional activity intensity (Zang et al., 2007), was computed at voxel level within range 0.01 – 0.1 Hz and then z-score transformation was applied to origin ALFF and averaged in all 246 ROIs for further analysis. Degree centrality (DC) was computed by summing all reserved structural probabilistic connectivity and functional connectivity for each ROI, representing regional structural DC (sDC) and functional DC (fDC) respectively.

### 2.7. Estimation of Relationship between Music Perception and Neuroimaging Features

A machine learning algorithm called L2-regularized L2-loss support vector regression (SVR) and a 10-fold cross validation method were employed to explore the relationship between averaged MBEA score and ALFF, sDC, fDC and structure-function coupling respectively (Figure 1A). We filtered features (Spearman correlation between MBEA score and structure-function coupling, uncorrected *p* < 0.05) and generated SVR model with default parameters using random 9 folds of all subjects, then predicted MBEA score in remaining 1-fold using features and model proposed by training folds. The correlation coefficient, mean absolute error (MAE) and mean square error (MSE) between actual and predicted MBEA score was calculated to evaluate prediction performance. A permutation test (5000 permutations) was performed to examine the significance of model. The contribution of each ROI of each feature was calculated by summing up their contributions in each cross validation (if a feature was not selected in one cross validation, its contribution to this cross validation was set to zero). Only ROIs selected in all cross validation and any predictive model were included in further analysis and discussion.

**Figure 1.**
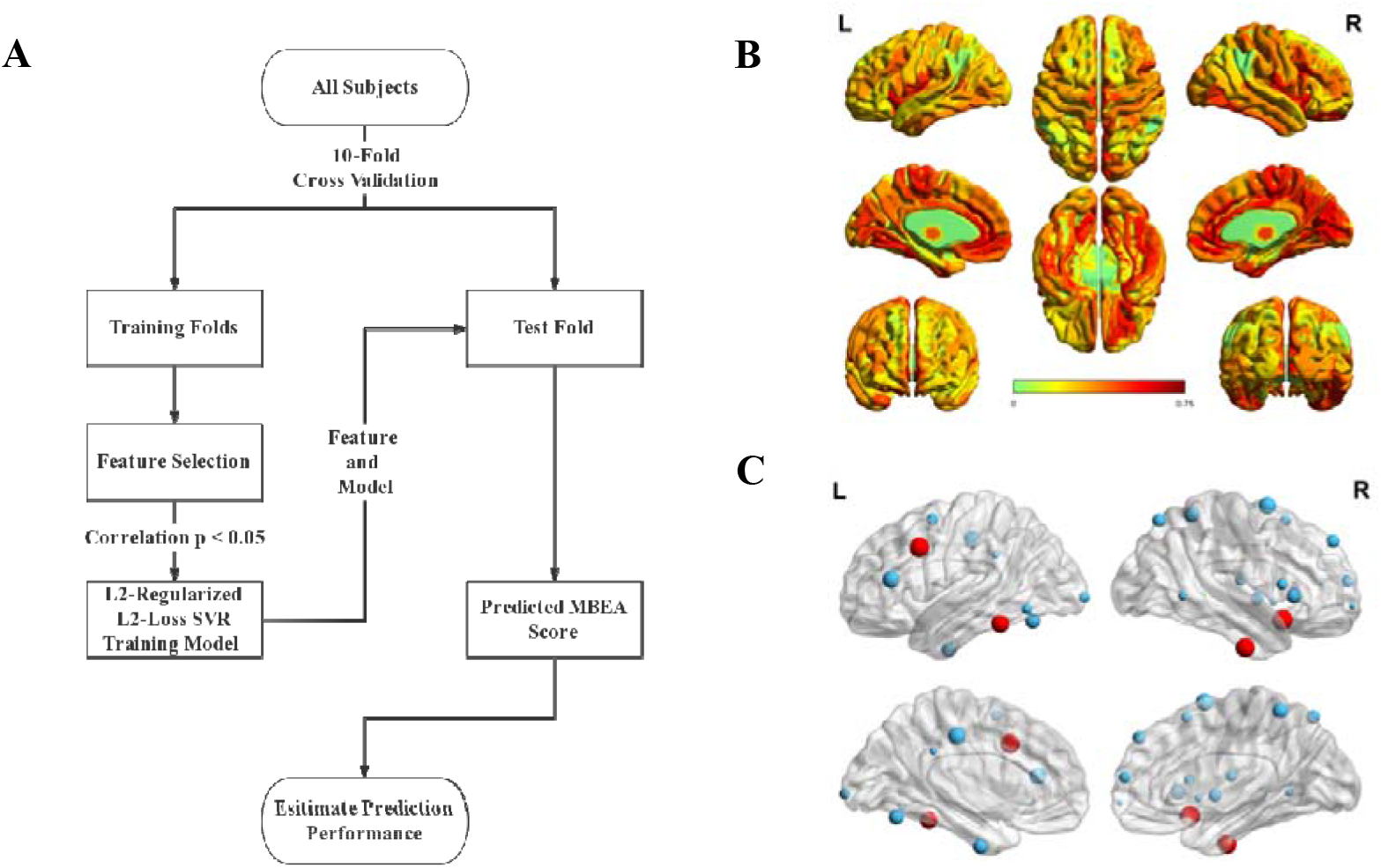
Prediction procedure, spatial pattern of mean structure-function coupling and all predictive ROIs. **(A)** Illustration of 10-fold cross validation in our study. In training folds, we filtered potential features (ALFF, functional or structural degree centrality or structure-function coupling) by Spearman correlation and trained prediction model. In test fold, we calculated predicted MBEA score using features and model from training folds. MAE, MSE and Spearman correlation were used to estimate prediction performance. **(B)** Hierarchical structure-function coupling across cortex. Primary and medial cortex showed relatively high, while higher-order and lateral cortex showed relatively low structure-function coupling. **(C)** Predictive ROIs distributed from subcortical to cortical areas. Greater volume represents greater contribution in prediction for each ROI. Most effective ROIs are marked with red color.

### 2.8. Exploratory Analysis

Meditation analysis was employed to explore the potential relationship between multiple neuroimaging features and how these relationships and features (independent or meditating variables) affect music perception (MBEA score). We conducted meditation analysis using the PROCESS plug-in in SPSS. We conducted correlation analysis between ROIs features and some irrelevant variables (hearing threshold and IQ), in order to examine the specificity of our results.

### 2.9. Association between Meta-analytic Cognitive Functions and Predictive ROIs

To describe the potential cognitive functions and implications of predictive ROIs, we utilized the *Decoder* function in Neurosynth (Yarkoni et al., 2011) to obtain the spatial correlation between predictive ROIs with or without their connected regions and meta-analytic map of terms in database. We presented analysis results as word clouds.

Our main result considered gender, age and mean Jenkinson’s FD as covariates. We also conducted the procedure above without covariates for a validation analysis.

## 3. Results

### 3.1. Participant Characteristics

In 106 participants were between the ages of 18 – 24 years of age and averaged MBEA score ranged from 14.5 to 29.5. 43 individuals of them diagnosed with CA according to suggested MBEA cut-off value. We didn’t find any correlation between MBEA score and age, gender, education years, IQ and head motion (all *p* > 0.05).

### 3.2. Prediction Performance of Four Features for MBEA Score

Similar with other studies, we found a hierarchical variation of structure-function coupling from primary to high-order cortex (Figure 1B). Then a set of structure-function coupling features showed predictive effect for averaged MBEA score (correlation *r* = .336, *p* < .001, MAE = 3.487, MSE = 18.263; permutation test *p* < .005; Figure 2A & 2B). Generally, extensive areas including frontal, temporal, parietal, occipital, insula and basal ganglia in both hemispheres contributed to predict MBEA score (Table 1 and Figure 1C). In which, right intermediate lateral and left caudolateral area 20 of inferior temporal gyrus (R.ITG_A20il, L.ITG_A20cl), left inferior frontal junction of MFG (L.MFG_IFJ) and right ventral agranular insula (R.Ins_vIa) were selected in 10 cross validation folds and showed greatest contribution for prediction. SVR model based on ALFF and DC failed to predict MBEA score (*r* = -.036, -.184, .024 for ALFF, sDC and fDC respectively; all correlation *p* > .05; Figure 2C - 2E).

**Table 1.**
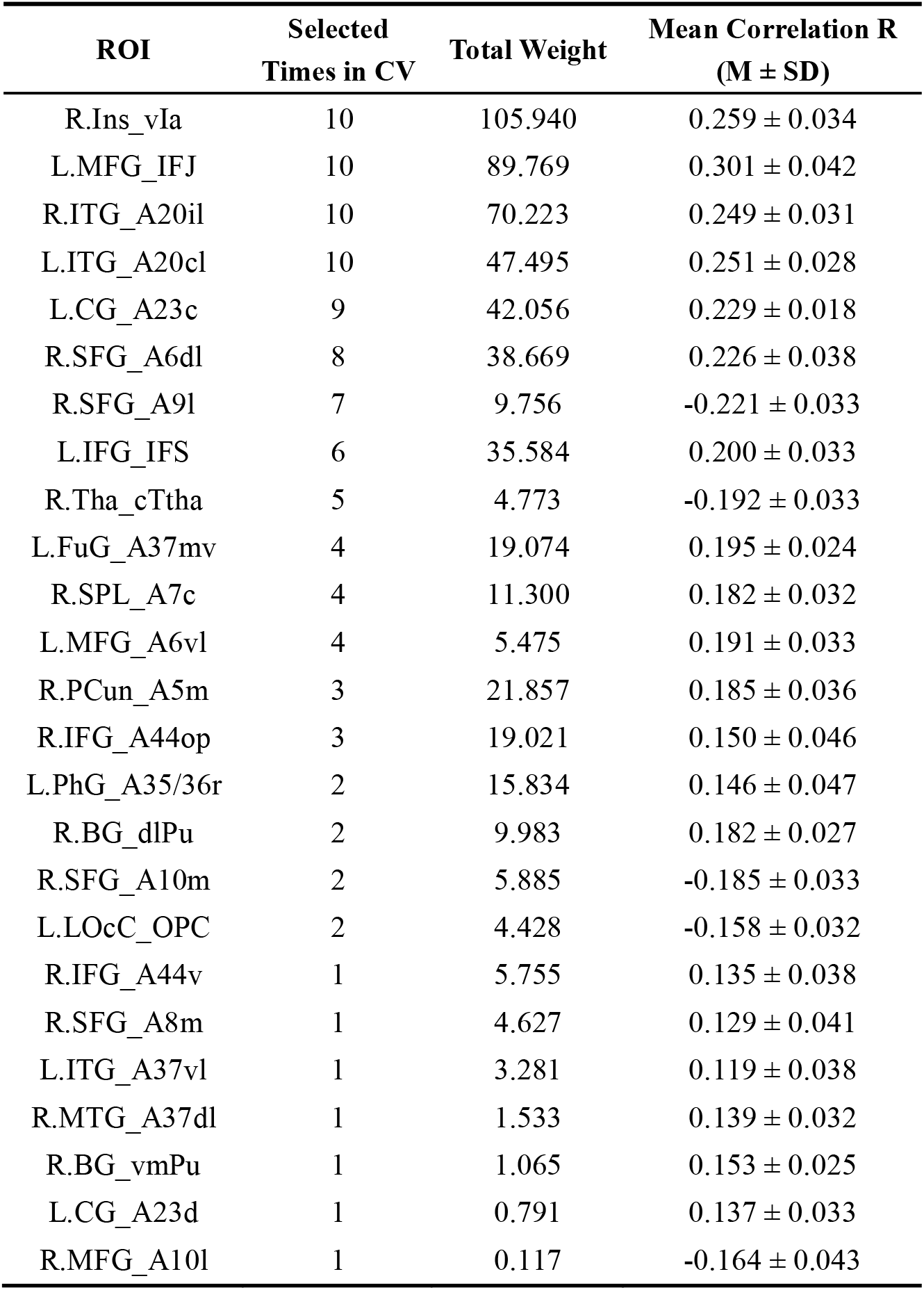
All predictive ROIs and their total weight, selected times and correlation coefficient across all 10 cross validations. **Abbreviations:** ROI: Region of Interest; M ± SD: Mean ± Standard Deviation; CV: Cross Validation; Ins: Insula; MFG: Middle Frontal Gyrus; ITG: Inferior Temporal Gyrus; CG: Cingulate Gyrus; SFG: Superior Frontal Gyrus; IFG: Inferior Frontal Gyrus; Tha: Thalamus; FuG: Fusiform Gyrus; SPL: Superior Parietal Lobule; PCun: Precuneus; PhG: Parahippocampus Gyrus; BG: Basal Ganglia; LOcC: lateral Occipital Cortex; MTG: Middle Temporal Gyrus.

**Figure 2.**
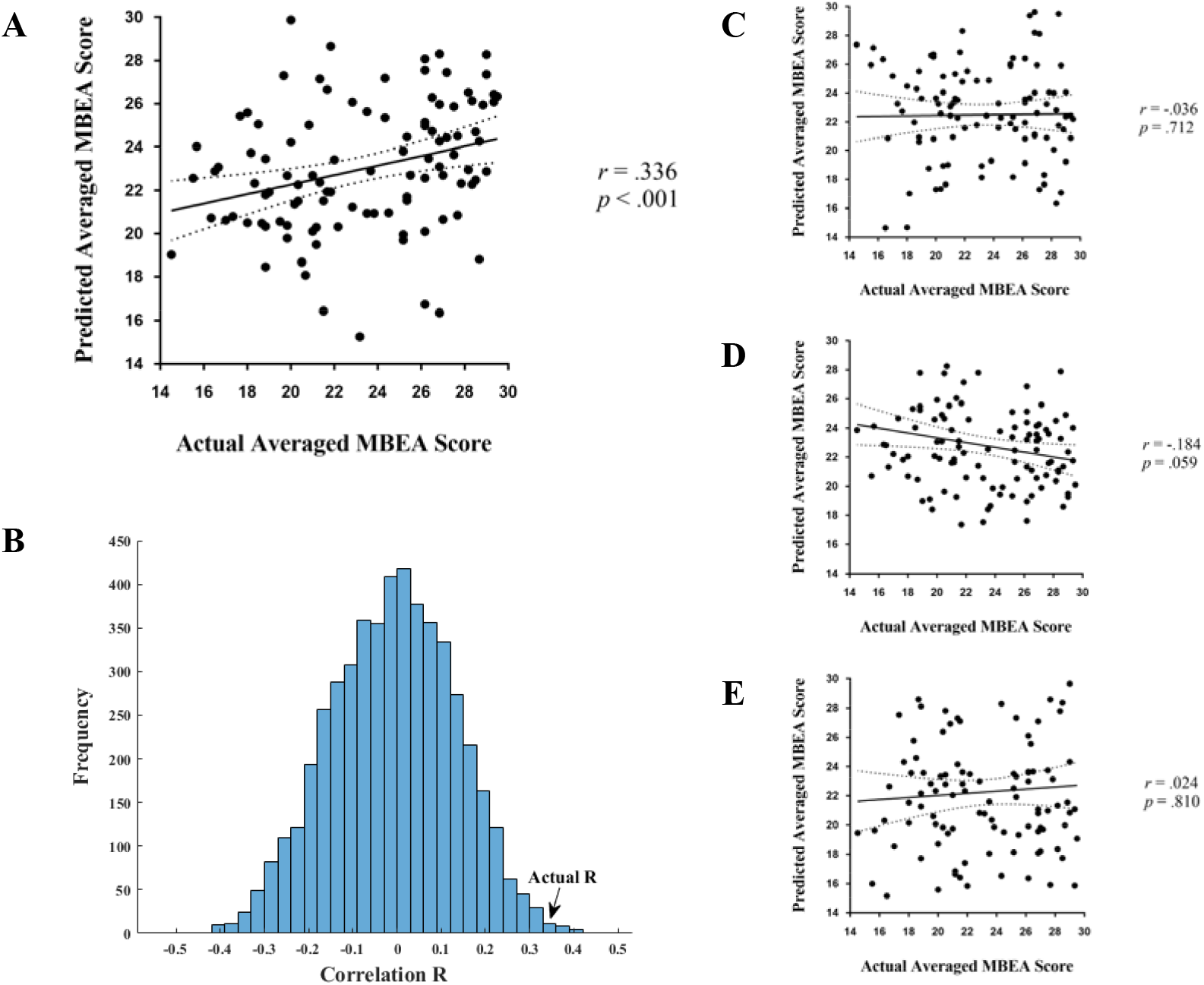
Performance of different feature-based prediction model. **(A)** Significant correlation between MBEA score predicted by structure-function coupling based model and actual MBEA score was found (correlation *r* = .336, *p* < .001). **(B)** Prediction performance of coupling based model was not accident according to permutation test (*p* < .005). **(C), (D) and (E)** Correlation between MBEA score predicted by ALFF, structural and functional degree centrality and actual MBEA score respectively. All these three features failed to predict MBEA score (all *p* > 0.05). Solid line and dashed lines in scatter plots represent best-fit line and 95% confidence interval.

All four most predictive ROIs mainly connected with frontotemporal areas of their ipsilateral hemisphere and also connected with other regions such as occipital, parietal, insula lobe and subcortical regions. Although in the contralateral hemisphere, L.MFG_IFJ showed a direct structural connectivity with right IPL which is a major node of right dorsal stream. Structural connectivity patterns of four regions are shown in Figure 3.

**Figure 3.**
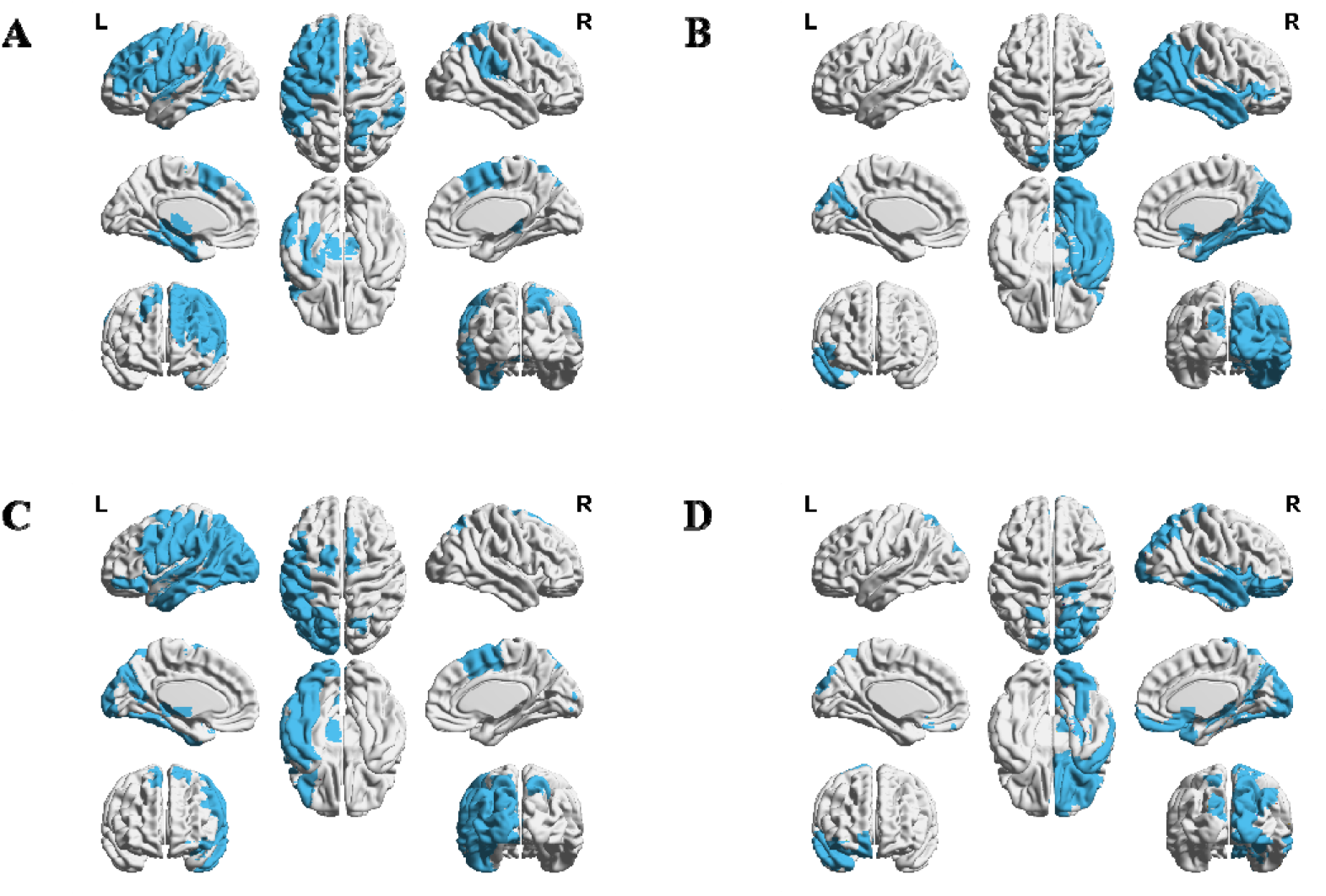
Structural connectivity patterns of most predictive four ROIs. **(A), (B), (C) and (D)** demonstrate connectivity patterns of L.MFG_IFJ, R.ITG_A20il, L.ITG_A20cl and R.Ins_vIa respectively. All four ROIs mainly connected with their ipsilateral frontotemporal areas, and also connected with occipital and parietal lobule. Particularly, L.MFG_IFJ showed a direct structural connectivity with contralateral IPL.

### 3.3. Mediated Effects and Specificity of Structure-function Coupling

Following the predictive effects of structure-function coupling on MBEA score reported above, we employed exploratory mediation analysis to explain the multiple relationship among structure-function coupling, sDC, fDC, ALFF, and MBEA score. We found a full indirect effect of mean ALFF (X variable) on music perception (MBEA score, Y variable) through mean structure-function coupling (M variable; partially standardized indirect effect = -.547, 95% CI: [-1.171, -.047]; Figure 4A) of most stable four ROIs. Separately, we captured this type of effect on L.MFG_IFJ (partially standardized indirect effect = -.323, 95% CI: [-.660, -.085]; Figure 4B).

**Figure 4.**
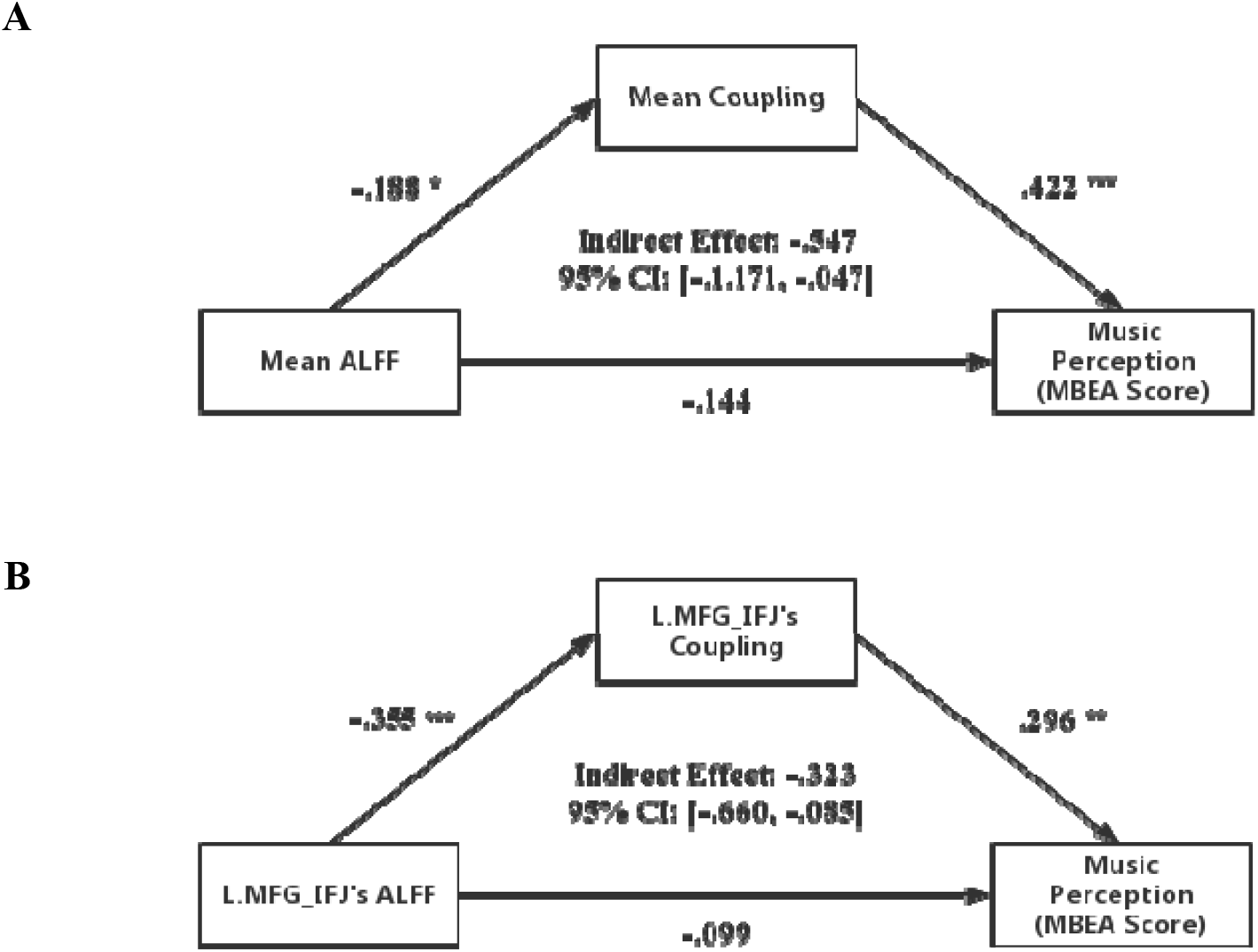
Mediated relation among ALFF, structure-function coupling and MBEA score. **(A)** Mean structure-function coupling of all four most predictive ROIs fully mediated the relation between mean ALFF and MBEA score (partially standardized indirect effect = -.547). **(B)** Structure-function coupling of L.MFG_IFJ fully mediated the relation between ALFF and MBEA score (partially standardized indirect effect = -.323).

We did not find any relation of structure-function coupling of most effective four ROIs to IQ and hearing threshold (all *p* > .05).

### 3.4. Meta-analytic Cognitive Functions Related to Predictive ROIs

The most predictive four ROIs were correlated with several meta-analytic cognitive terms such as language, phonological, semantic, memory, retrieval, judgement, cognitive/executive control. Because L.MFG_IFJ and L.ITG_A20cl showed some interesting cognitive-terms potentially associated with music perception and both locate out of widely accepted neural circuits underlying music processing. We demonstrated these cognitive terms in Figure 5. Cognitive terms of R.ITG_A20il and R.Ins_vIa were presented in supplementary materials (Table S2 & S3).

**Figure 5.**
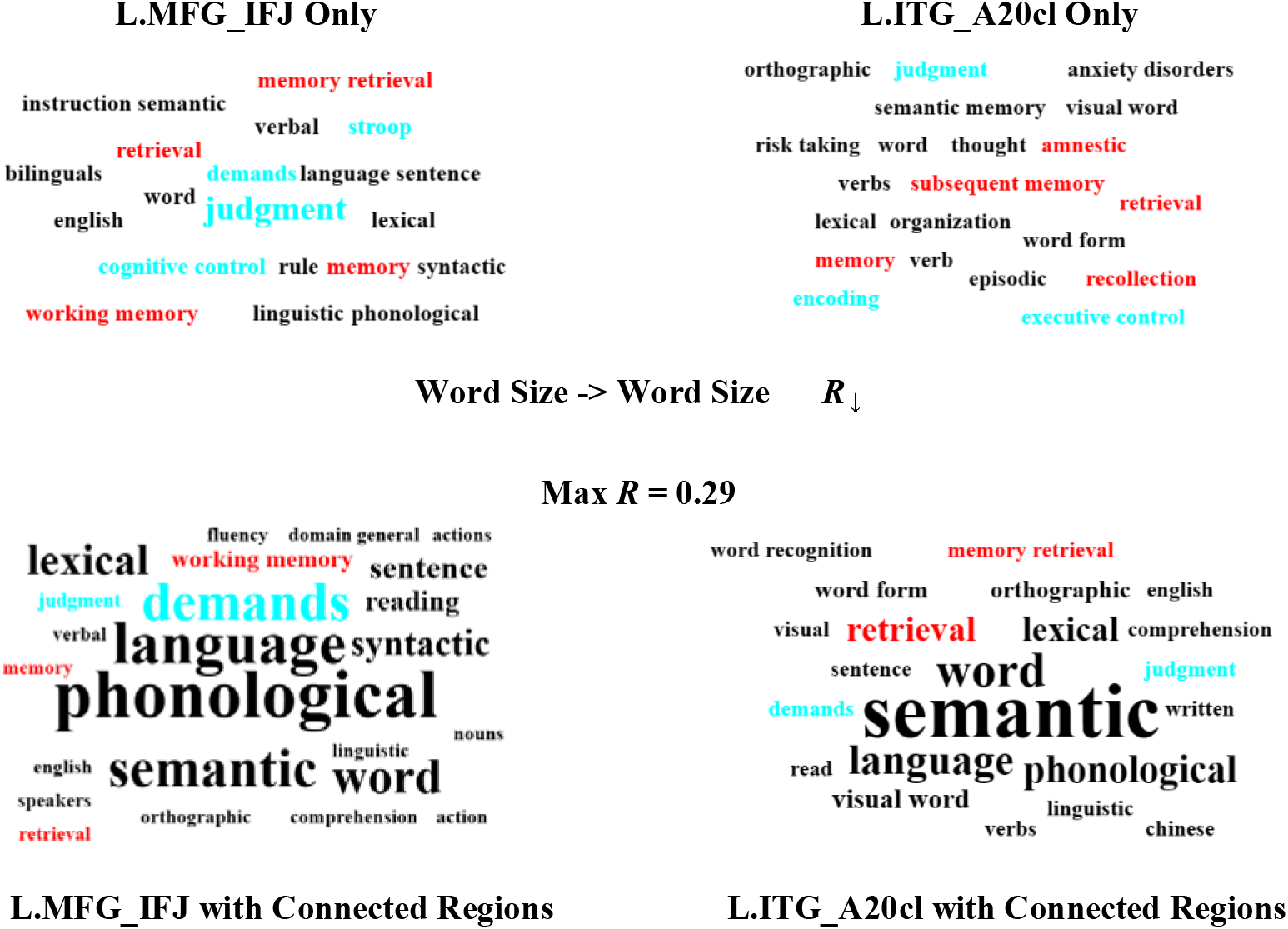
Cognitive meaning of L.MFG_IFJ and L.ITG_A20cl with or without their connected regions. Word clouds presented related cognitive functions. Greater word size represents stronger spatial correlations between regions and cognitive terms in Neurosynth (max correlation coefficient = 0.29). Some terms potentially associated with music perception were marked with non-black colors.

### 3.5. Validation Analysis

Except mediated effect of mean structure-function coupling of most predictive ROIs, similar results were found when we considered age, gender and head motion as covariates or not. Results of validation analysis were shown in supplementary materials (Figure S1 – S3, Table S1).

## 4. Discussion

Understanding the neural basis of music processing is an important issue of cognitive neuroscience. In this study, we investigated music related structure-function coupling, ALFF and sDC and fDC, and explored how does these features influence music perception. Instead of ALFF and both DC, we found widely distributed structure-function of brain regions from cortical to subcortical predicted music perception ability. Structure-function coupling of L.MFG_IFJ, L.ITG_A20cl, R.ITG_A20il and R.Ins_vIa made most effective contributions for prediction. Mean structure-function coupling of four regions above, especially L.MFG_IFJ, fully mediated the relation between their ALFF and music perception.

Right insula is a key node of structural pathway between IFG and STG, previous studies indicated insula engaged in multiple musical cognition (Putkinen et al., 2021; Van’t Hooft et al., 2021; Vuong et al., 2023) and altered structure or function in right insula might disturb the neural activity of right ventral stream of music neural processing. Indeed, lesion areas converged on right insula were reported in acquired amusia (Sihvonen et al., 2019). We found decreasing structure-function coupling of R.Ins_vIa contributed to worse music perception, which is consistent with opinion above. ITG is another brain region of frontotemporal network and is connected with STG (Lin et al., 2020). Altered structure and function of ITG involved in amusia according to previous researches (Sihvonen et al., 2019). We also found a positive relationship between structure-function coupling of R.ITG_A20il and music perception. These results provide further support for the dual-stream model of music perception.

We found structure-function coupling of wide brain regions is predictive, but ALFF and both DC failed to explain music perception. Structure-function coupling is implicit in structural constraint, maintenance and modulation of functional communication in certain brain regions or circuits (Avena-Koenigsberger et al., 2017; Gomez et al., 2019; Liu et al., 2022). These results indicated that uncoordinated neural activity due to decoupled structure and function, but not regional activity or connectivity underlies music perception. High plasticity of human brain and/or interaction between structure and function might bring unpredictable changes either in structure or function across lifespan, which increases the difficulty and lead to failure when exploring brain-behavior association via single modal neuroimaging features. Structure-function coupling could capture this gap, therefore structure-function coupling is predictive for music perception in our study and might be more effective for understanding other cognitive functions. Moreover, we found mean structure-function coupling of most predictive ROIs fully mediated the relation between ALFF and music perception, which further explain the reason why regional activity failed to predict music perception: elevated spontaneous regional neural activity leads to poor music perception by breaking their communications between brain regions in way of disassociating structure and function.

Particularly, we found a fully negative indirect effect of L.MFG_IFS’s ALFF on music perception. L.MFG_IFS is structurally connected with right neural circuit related with music perception, our results were inconsistent with the classical view which considered that upregulating adaptive changes of contralateral hemispheric homologous areas of certain function regions play a compensatory role for their insufficient efficacy (Bettus et al., 2009; Blesa et al., 2011; Knights et al., 2021), especially in lesion-brains. Some recent studies challenged this opinion and argued that the up-regulation in contralateral hemisphere might be maladaptive (Stefaniak et al., 2020). Keser et al indicated that poor language recovery after stroke was associated with higher tract integrity of right AF (Keser et al., 2020). More causally, Ren et al suggested that inhibitory transcranial magnetic stimulation (TMS) targeted in right contributed to recovery of aphasia and demonstrated comparable treatment efficacy to excitatory TMS therapy targeted in left superior frontal gyrus (Ren et al., 2023). Aphasia is a disease with similar neural mechanism and pattern of symptoms to amusia, these findings are of certain reference value for music and amusia. Increased inter-hemisphere connectivity was also reported in amusia (Albouy et al., 2013; Jin et al., 2021a). MFG_IFS is adjacent to IFG, and it was demonstrated right dorsolateral prefrontal cortex engages in, even provides causal support for music perception (Albouy et al., 2018; Chen and Yuan, 2016). Combined with these findings, our results indicated elevated activity on left hemispheric homologous music circuits is a maladaptive regulation. Inhibitory neuromodulation therapy targeted in contralateral homologous regions might be useful for post-brain damage amusia because of both improving effect of traditional treatment and enhance tolerance of therapy targeted in ipsilateral music related areas.

We can draw some inferences about cognitive causes of amusia according to our results and cognitive neuroscience knowledge. Anterior insula (aIns) is the integral core hub of salience network, which assists target brain regions to generate appropriate cognitive responses and guide behavior to salient stimuli (Menon and Uddin, 2010). When salient stimulations are detected, aIns participate in modulating autonomic reactivity and facilitating access to attention and working memory resources. Meanwhile, we found altered structure-function coupling of R.Ins_vIa, L.MFG_IFS and L.ITG_A20cl underlying music perception and the latter two regions involved in cognitive functions such as memory (working memory, memory retrieval, amnestic), cognitive control and judgement according to Neurosynth. Therefore, we argue that abnormal attention, and/or abnormal memory storage and/or retrieval for music stimulations may be potential cognitive causes of amusia, which agree with neurobiology framework of auditory learning (Kraus and White-Schwoch, 2015) and amusia (Peretz, 2016).

There are several limitations of present study. First, the small sample size of our study and lack of independent test data set limits the generalizability of our findings. Second, we found predictive contributions of regions in both hemispheres are substantially the same, which disagree with right lateralization of musical neural processing. Pitkäniemi et al suggested singing music lean on left lateralized pathways (Pitkäniemi et al., 2023). Neural lateralization of music needs further investigation. In addition, we gave some “brave” assumptions about cognitive causes of music processing although they had been proven to a certain extent (Albouy et al., 2013; Albouy et al., 2015; Albouy et al., 2018). All these findings and assumptions needs further exploration and validation.

## 5. Conclusions

Our findings indicate that structure-function coupling is more effective when capturing neural correlates of music, and increased activity of contralateral regions homologous with music pathways account for worse music perception via decoupling the association between structure and function. Altogether, our findings propose a new perspective to neural basis of music and plasticity, and offer insights to neuromodulation therapy and psychological processing of music.

## Ethics statement

Our study was approved by Ethics Committee of the Second Xiangya Hospital of Central South University. All participants provided informed consent through their signatures.

## Data availability

All data that support these findings of current study are available on request from the corresponding author, upon reasonable request.

## CRediT authorship contribution statement

Yuchen Wang: Data curation, Formal analysis, Investigation, Visualization, Writing – original draft. Zhishuai Jin: Investigation, Writing – review & editing. Sizhu Huyang: Investigation, Writing – review & editing. Qiaoping Lian: Investigation, Writing – review & editing. Daxing Wu: Conceptualization, Funding acquisition, Supervision, Data curation, Writing – review & editing, Resources, Project administration.

## Acknowledgments

This work was supported by research grants from National Natural Science Foundation of China awarded to D.W. (No. 81771172).

## References

Albouy, P., Caclin, A., Norman-Haignere, S.V., Lévêque, Y., Peretz, I., Tillmann, B., Zatorre, R.J., 2019. Decoding Task-Related Functional Brain Imaging Data to Identify Developmental Disorders: The Case of Congenital Amusia. Frontiers in Neuroscience 13.

Albouy, P., Mattout, J., Bouet, R., Maby, E., Sanchez, G., Aguera, P.E., Daligault, S., Delpuech, C., Bertrand, O., Caclin, A., Tillmann, B., 2013. Impaired pitch perception and memory in congenital amusia: the deficit starts in the auditory cortex. Brain 136, 1639–1661.

Albouy, P., Mattout, J.r.m., Sanchez, G.t., Tillmann, B., Caclin, A., 2015. Altered retrieval of melodic information in congenital amusia: insights from dynamic causal modeling of MEG data. Frontiers in Human Neuroscience 9.

Albouy, P., Peretz, I., Bermudez, P., Zatorre, R.J., Tillmann, B., Caclin, A., 2018. Specialized neural dynamics for verbal and tonal memory: fMRI evidence in congenital amusia. Human Brain Mapping 40, 855–867.

Avena-Koenigsberger, A., Misic, B., Sporns, O., 2017. Communication dynamics in complex brain networks. Nature Reviews: Neuroscience 19, 17–33.

Baum, G.L., Cui, Z., Roalf, D.R., Ciric, R., Betzel, R.F., Larsen, B., Cieslak, M., Cook, P.A., Xia, C.H., Moore, T.M., Ruparel, K., Oathes, D.J., Alexander-Bloch, A.F., Shinohara, R.T., Raznahan, A., Gur, R.E., Gur, R.C., Bassett, D.S., Satterthwaite, T.D., 2019. Development of structure–function coupling in human brain networks during youth. Proceedings of the National Academy of Sciences 117, 771–778.

Bettus, G., Guedj, E., Joyeux, F., Confort-Gouny, S., Soulier, E., Laguitton, V., Cozzone, P.J., Chauvel, P., Ranjeva, J.P., Bartolomei, F., Guye, M., 2009. Decreased basal fMRI functional connectivity in epileptogenic networks and contralateral compensatory mechanisms. Human Brain Mapping 30, 1580–1591.

Blesa, J., Juri, C., García-Cabezas, M., Adánez, R., Sánchez-González, M., Cavada, C., Obeso, J.A., 2011. Inter-hemispheric asymmetry of nigrostriatal dopaminergic lesion: a possible compensatory mechanism in Parkinson’s disease. Frontiers in Systems Neuroscience 5, 92.

Buchanan, C.R., Bastin, M.E., Ritchie, S.J., Liewald, D.C., Madole, J.W., Tucker-Drob, E.M., Deary, I.J., Cox, S.R., 2020. The effect of network thresholding and weighting on structural brain networks in the UK Biobank. Neuroimage 211.

Cao, R., Wang, X., Gao, Y., Li, T., Zhang, H., Hussain, W., Xie, Y., Wang, J., Wang, B., Xiang, J., 2020. Abnormal Anatomical Rich-Club Organization and Structural–Functional Coupling in Mild Cognitive Impairment and Alzheimer’s Disease. Frontiers in Neurology 11.

Chan, S.Y., Ong, Z.Y., Ngoh, Z.M., Chong, Y.S., Zhou, J.H., Fortier, M.V., Daniel, L.M., Qiu, A., Meaney, M.J., Tan, A.P., 2022. Structure-function coupling within the reward network in preschool children predicts executive functioning in later childhood. Developmental Cognitive Neuroscience 55.

Chen, J., Yuan, J., 2016. The Neural Causes of Congenital Amusia. Journal of Neuroscience 36, 7803–7804.

Chen, Q., Lv, H., Wang, Z., Wei, X., Liu, J., Liu, F., Zhao, P., Yang, Z., Gong, S., Wang, Z., 2022. Distinct brain structuralLfunctional network topological coupling explains different outcomes in tinnitus patients treated with sound therapy. Human Brain Mapping 43, 3245–3256.

Collin, G., Scholtens, L.H., Kahn, R.S., Hillegers, M.H.J., van den Heuvel, M.P., 2017. Affected Anatomical Rich Club and Structural–Functional Coupling in Young Offspring of Schizophrenia and Bipolar Disorder Patients. Biological Psychiatry 82, 746–755.

Fan, L., Li, H., Zhuo, J., Zhang, Y., Wang, J., Chen, L., Yang, Z., Chu, C., Xie, S., Laird, A.R., Fox, P.T., Eickhoff, S.B., Yu, C., Jiang, T., 2016. The Human Brainnetome Atlas: A New Brain Atlas Based on Connectional Architecture. Cerebral Cortex 26, 3508–3526.

Genon, S., Reid, A., Langner, R., Amunts, K., Eickhoff, S.B., 2018. How to Characterize the Function of a Brain Region. Trends in Cognitive Sciences 22, 350–364.

Gomez, J., Drain, A., Jeska, B., Natu, V.S., Barnett, M., Grill-Spector, K., 2019. Development of population receptive fields in the lateral visual stream improves spatial coding amid stable structural-functional coupling. Neuroimage 188, 59–69.

Gu, Z., Jamison, K.W., Sabuncu, M.R., Kuceyeski, A., 2021. Heritability and interindividual variability of regional structure-function coupling. Nature Communications 12.

Hyde, K.L., Lerch, J.P., Zatorre, R.J., Griffiths, T.D., Evans, A.C., Peretz, I., 2007. Cortical thickness in congenital amusia: when less is better than more. Journal of Neuroscience 27, 13028–13032.

Jin, Z., Huyang, S., Jiang, L., Yan, Y., Xu, M., Wang, J., Li, Q., Wu, D., 2021a. Increased Resting-State Interhemispheric Functional Connectivity of Posterior Superior Temporal Gyrus and Posterior Cingulate Cortex in Congenital Amusia. Frontiers in Neuroscience 15.

Jin, Z., Lu, X., Huyang, S., Yan, Y., Jiang, L., Wang, J., Xu, M., Li, Q., Wu, D., 2021b. Impaired face recognition is associated with abnormal gray matter volume in the posterior cingulate cortex in congenital amusia. Neuropsychologia 156.

Keser, Z., Sebastian, R., Hasan, K.M., Hillis, A.E., 2020. Right Hemispheric Homologous Language Pathways Negatively Predicts Poststroke Naming Recovery. Stroke 51, 1002–1005.

Knights, E., Morcom, A.M., Henson, R.N., 2021. Does Hemispheric Asymmetry Reduction in Older Adults in Motor Cortex Reflect Compensation? Journal of Neuroscience 41, 9361–9373.

Koubiyr, I., Besson, P., Deloire, M., Charre-Morin, J., Saubusse, A., Tourdias, T., Brochet, B., Ruet, A., 2019. Dynamic modular-level alterations of structural-functional coupling in clinically isolated syndrome. Brain 142, 3428–3439.

Kraus, N., White-Schwoch, T., 2015. Unraveling the Biology of Auditory Learning: A Cognitive-Sensorimotor-Reward Framework. Trends in Cognitive Sciences 19, 642–654.

Kulik, S.D., Nauta, I.M., Tewarie, P., Koubiyr, I., van Dellen, E., Ruet, A., Meijer, K.A., de Jong, B.A., Stam, C.J., Hillebrand, A., Geurts, J.J.G., Douw, L., Schoonheim, M.M., 2022. Structure-function coupling as a correlate and potential biomarker of cognitive impairment in multiple sclerosis. Network Neuroscience 6, 339–356.

Liao, X., Sun, J., Jin, Z., Wu, D., Liu, J., 2022. Cortical Morphological Changes in Congenital Amusia: Surface-Based Analyses. Frontiers in Psychiatry 12.

Lin, Y.H., Young, I.M., Conner, A.K., Glenn, C.A., Chakraborty, A.R., Nix, C.E., Bai, M.Y., Dhanaraj, V., Fonseka, R.D., Hormovas, J., Tanglay, O., Briggs, R.G., Sughrue, M.E., 2020. Anatomy and White Matter Connections of the Inferior Temporal Gyrus. World Neurosurgery 143, e656–e666.

Litwińczuk, M.C., Muhlert, N., Cloutman, L., Trujillo-Barreto, N., Woollams, A., 2022. Combination of structural and functional connectivity explains unique variation in specific domains of cognitive function. Neuroimage 262, 119531.

Liu, X., Qiu, S., Wang, X., Chen, H., Tang, Y., Qin, Y., 2023a. Aberrant dynamic Functional-Structural connectivity coupling of Large-scale brain networks in poststroke motor dysfunction. NeuroImage: Clinical 37.

Liu, Z.-Q., Shafiei, G., Baillet, S., Misic, B., 2023b. Spatially heterogeneous structure-function coupling in haemodynamic and electromagnetic brain networks. Neuroimage 278.

Liu, Z.-Q., Vázquez-Rodríguez, B., Spreng, R.N., Bernhardt, B.C., Betzel, R.F., Misic, B., 2022. Time-resolved structure-function coupling in brain networks. Communications Biology 5.

Mas-Herrero, E., Singer, N., Ferreri, L., McPhee, M., Zatorre, R.J., Ripollés, P., 2023. Music engagement is negatively correlated with depressive symptoms during the COVID-19 pandemic via reward-related mechanisms. Annals of the New York Academy of Sciences 1519, 186–198.

Menon, V., Uddin, L.Q., 2010. Saliency, switching, attention and control: a network model of insula function. Brain Structure & Function 214, 655–667.

Nan, Y., Sun, Y., Peretz, I., 2010. Congenital amusia in speakers of a tone language: association with lexical tone agnosia. Brain 133, 2635–2642.

Norman-Haignere, S.V., Albouy, P., Caclin, A., McDermott, J.H., Kanwisher, N.G., Tillmann, B., 2016. Pitch-Responsive Cortical Regions in Congenital Amusia. Journal of Neuroscience 36, 2986–2994.

Pan, Y., Li, X., Liu, Y., Jia, X., Wang, S., Ji, Q., Zhao, W., Yin, B., Bai, G., Zhang, J., Bai, L., 2023. Hierarchical brain structural–functional coupling associated with cognitive impairments in mild traumatic brain injury. Cerebral Cortex 33, 7477–7488.

Peretz, I., 2016. Neurobiology of Congenital Amusia. Trends in Cognitive Sciences 20, 857–867.

Peretz, I., Champod, A.S., Hyde, K., 2003. Varieties of musical disorders. The Montreal Battery of Evaluation of Amusia. Annals of the New York Academy of Sciences 999, 58–75.

Pitkäniemi, A., Särkämö, T., Siponkoski, S.-T., Brownsett, S.L.E., Copland, D.A., Sairanen, V., Sihvonen, A.J., 2023. Hodological organization of spoken language production and singing in the human brain. Communications Biology 6.

Putkinen, V., Nazari-Farsani, S., Seppälä, K., Karjalainen, T., Sun, L., Karlsson, H.K., Hudson, M., Heikkilä, T.T., Hirvonen, J., Nummenmaa, L., 2021. Decoding Music-Evoked Emotions in the Auditory and Motor Cortex. Cerebral Cortex 31, 2549–2560.

Ren, J., Ren, W., Zhou, Y., Dahmani, L., Duan, X., Fu, X., Wang, Y., Pan, R., Zhao, J., Zhang, P., Wang, B., Yu, W., Chen, Z., Zhang, X., Sun, J., Ding, M., Huang, J., Xu, L., Li, S., Wang, W., Xie, W., Zhang, H., Liu, H., 2023. Personalized functional imaging-guided rTMS on the superior frontal gyrus for post-stroke aphasia: A randomized sham-controlled trial. Brain Stimulation 16, 1313–1321.

Roberts, J.A., Perry, A., Roberts, G., Mitchell, P.B., Breakspear, M., 2017. Consistency-based thresholding of the human connectome. Neuroimage 145, 118–129.

Sihvonen, A.J., Ripollés, P., Leo, V., Rodríguez-Fornells, A., Soinila, S., Särkämö, T., 2016. Neural Basis of Acquired Amusia and Its Recovery after Stroke. Journal of Neuroscience 36, 8872–8881.

Sihvonen, A.J., Ripollés, P., Rodríguez-Fornells, A., Soinila, S., Särkämö, T., 2017a. Revisiting the Neural Basis of Acquired Amusia: Lesion Patterns and Structural Changes Underlying Amusia Recovery. Frontiers in Neuroscience 11.

Sihvonen, A.J., Sammler, D., Ripollés, P., Leo, V., RodríguezLFornells, A., Soinila, S., Särkämö, T., 2021. Right ventral stream damage underlies both poststroke aprosodia and amusia. European Journal of Neurology 29, 873–882.

Sihvonen, A.J., Särkämö, T., Ripollés, P., Leo, V., Saunavaara, J., Parkkola, R., Rodríguez-Fornells, A., Soinila, S., 2017b. Functional neural changes associated with acquired amusia across different stages of recovery after stroke. Scientific Report 7, 11390.

Sihvonen, A.J., Särkämö, T., Rodríguez-Fornells, A., Ripollés, P., Münte, T.F., Soinila, S., 2019. Neural architectures of music – Insights from acquired amusia. Neuroscience & Biobehavioral Reviews 107, 104–114.

Stefaniak, J.D., Halai, A.D., Lambon Ralph, M.A., 2020. The neural and neurocomputational bases of recovery from post-stroke aphasia. Nature Reviews: Neuroscience 16, 43–55.

Sun, J.-J., Pan, X.-Q., Yang, R., Jin, Z.-S., Li, Y.-H., Wu, D.-X., Liu, J., 2021. Changes in sensorimotor regions of the cerebral cortex in congenital amusia: a case-control study. Neural Regeneration Research 16.

Tay, J., Düring, M., van Leijsen, E.M.C., Bergkamp, M.I., Norris, D.G., de Leeuw, F.-E., Markus, H.S., Tuladhar, A.M., 2023. Network structure-function coupling and neurocognition in cerebral small vessel disease. NeuroImage: Clinical 38.

Tononi, G., Sporns, O., Edelman, G.M., 1994. A measure for brain complexity: relating functional segregation and integration in the nervous system. Proceedings of the National Academy of Sciences 91, 5033–5037.

Valk, S.L., Xu, T., Paquola, C., Park, B.-y., Bethlehem, R.A.I., Vos de Wael, R., Royer, J., Masouleh, S.K., Bayrak, ş., Kochunov, P., Yeo, B.T.T., Margulies, D., Smallwood, J., Eickhoff, S.B., Bernhardt, B.C., 2022. Genetic and phylogenetic uncoupling of structure and function in human transmodal cortex. Nature Communications 13.

Van’t Hooft, J.J., Pijnenburg, Y.A.L., Sikkes, S.A.M., Scheltens, P., Spikman, J.M., Jaschke, A.C., Warren, J.D., Tijms, B.M., 2021. Frontotemporal dementia, music perception and social cognition share neurobiological circuits: A meta-analysis. Brain and Cognition 148, 105660.

Vázquez-Rodríguez, B., Suárez, L.E., Markello, R.D., Shafiei, G., Paquola, C., Hagmann, P., van den Heuvel, M.P., Bernhardt, B.C., Spreng, R.N., Misic, B., 2019. Gradients of structure–function tethering across neocortex. Proceedings of the National Academy of Sciences 116, 21219–21227.

Vuong, V., Hewan, P., Perron, M., Thaut, M.H., Alain, C., 2023. The neural bases of familiar music listening in healthy individuals: An activation likelihood estimation meta-analysis. Neuroscience & Biobehavioral Reviews 154, 105423.

Wang, J., Khosrowabadi, R., Ng, K.K., Hong, Z., Chong, J.S.X., Wang, Y., Chen, C.-Y., Hilal, S., Venketasubramanian, N., Wong, T.Y., Chen, C.L.-H., Ikram, M.K., Zhou, J., 2018. Alterations in Brain Network Topology and Structural-Functional Connectome Coupling Relate to Cognitive Impairment. Frontiers in Aging Neuroscience 10.

Wu, D., Wang, X., Lin, S., Xu, G., Tian, J., Ma, X., 2023. Predicting insomnia severity using structure-function coupling in female chronic insomnia patients. Behavioural Brain Research 441.

Yan, C.G., Wang, X.D., Zuo, X.N., Zang, Y.F., 2016. DPABI: Data Processing & Analysis for (Resting-State) Brain Imaging. Neuroinformatics 14, 339–351.

Yarkoni, T., Poldrack, R.A., Nichols, T.E., Van Essen, D.C., Wager, T.D., 2011. Large-scale automated synthesis of human functional neuroimaging data. Nature Methods 8, 665–670.

Zang, Y.F., He, Y., Zhu, C.Z., Cao, Q.J., Sui, M.Q., Liang, M., Tian, L.X., Jiang, T.Z., Wang, Y.F., 2007. Altered baseline brain activity in children with ADHD revealed by resting-state functional MRI. Brain and Development 29, 83–91.

Zarkali, A., McColgan, P., Leyland, L.-A., Lees, A.J., Rees, G., Weil, R.S., 2021. Organisational and neuromodulatory underpinnings of structural-functional connectivity decoupling in patients with Parkinson’s disease. Communications Biology 4.

Zhang, R., Shao, R., Xu, G., Lu, W., Zheng, W., Miao, Q., Chen, K., Gao, Y., Bi, Y., Guan, L., McIntyre, R.S., Deng, Y., Huang, X., So, K.F., Lin, K., 2019. Aberrant brain structural–functional connectivity coupling in euthymic bipolar disorder. Human Brain Mapping 40, 3452–3463.

Zhang, S., Xu, X., Li, Q., Chen, J., Liu, S., Zhao, W., Cai, H., Zhu, J., Yu, Y., 2022. Brain Network Topology and Structural–Functional Connectivity Coupling Mediate the Association Between Gut Microbiota and Cognition. Frontiers in Neuroscience 16.

Zhang, Z., Liao, W., Chen, H., Mantini, D., Ding, J.R., Xu, Q., Wang, Z., Yuan, C., Chen, G., Jiao, Q., Lu, G., 2011. Altered functional-structural coupling of large-scale brain networks in idiopathic generalized epilepsy. Brain 134, 2912–2928.

