## Supplemental Figure for "Elevated Activity in Left Homologous Music Circuits is Maladaptive for Music Perception but Mediated by Decoupled Structure and Function"

**Figure S1. Performance of different feature-based prediction model without covariates.** Same with our main results, **(A)** Actual MBEA score could be predicted by structure-function coupling based model (correlation *r* = .241, *p* = .013; MAE = 3.749, MSE = 19.681). **(B)** Prediction performance of coupling based model was not accident according to permutation test (*p* = .035). **(C), (D) and (E)** Correlation between MBEA score predicted by ALFF, structural and functional degree centrality and actual MBEA score respectively. All these three features also failed to predict MBEA score (correlation *r* = .115, -.159, -.029 for ALFF, sDC and fDC respectively; all correlation *p* > .05). Solid line and dashed lines in scatter plots represent best-fit line and 95% confidence interval.

**A**

**B**

**C**

**D**

**E**

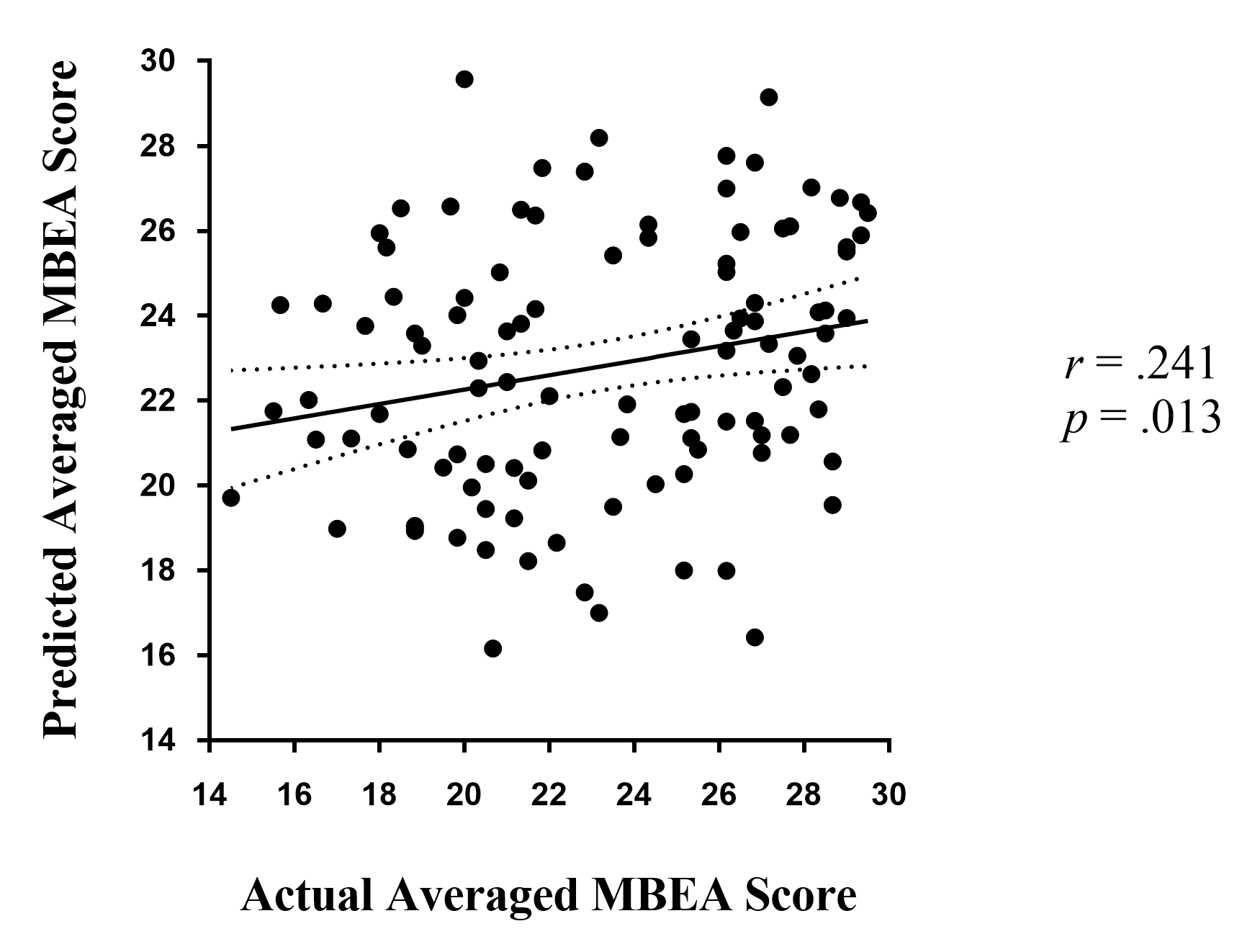

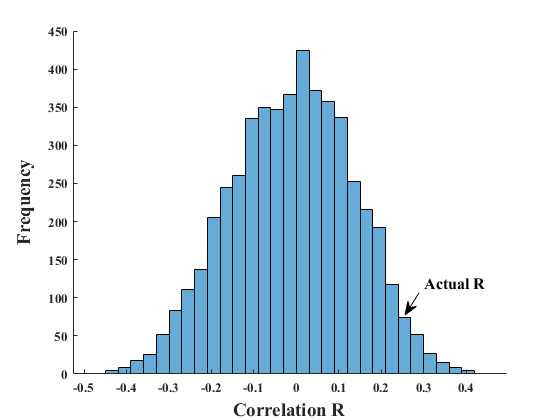

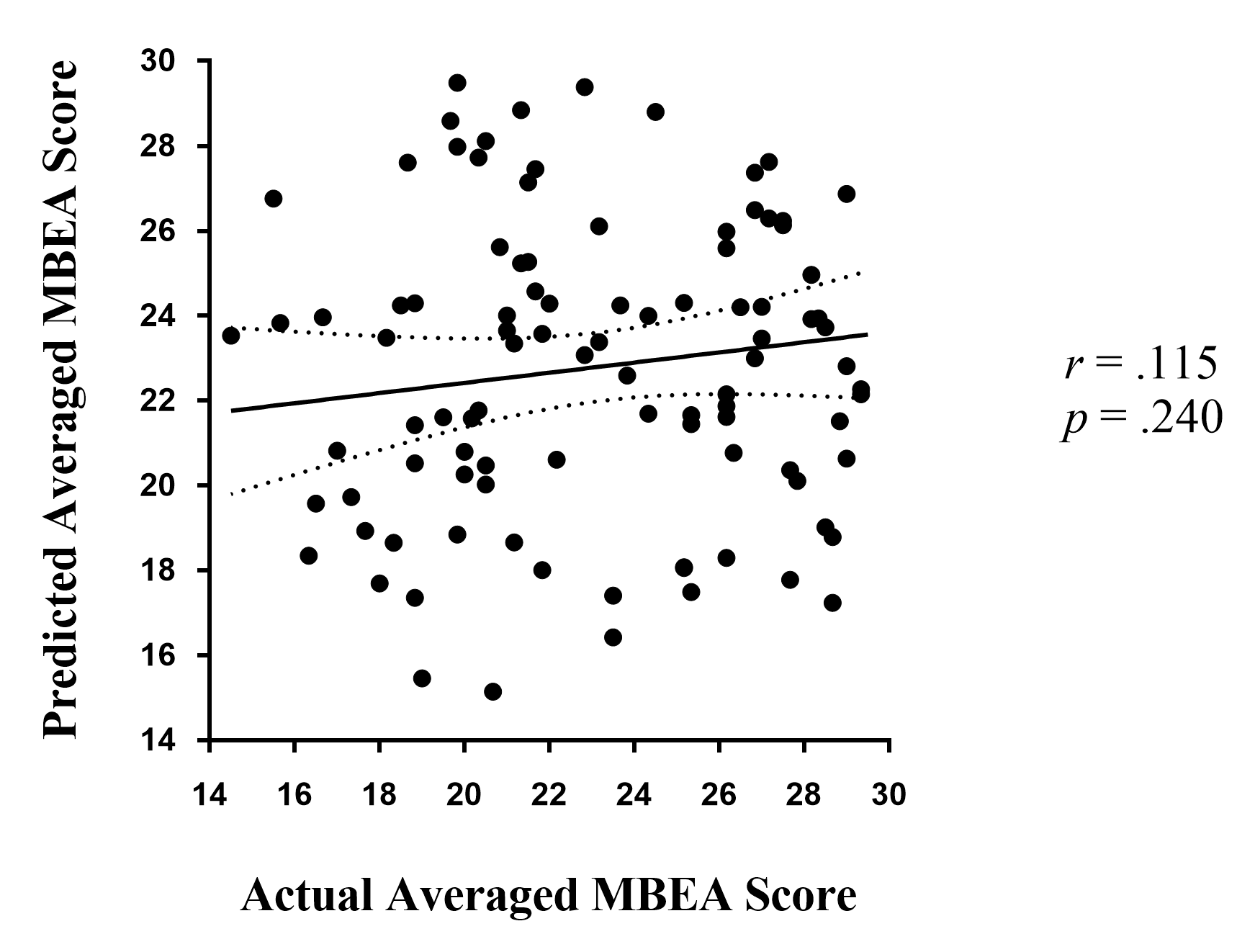

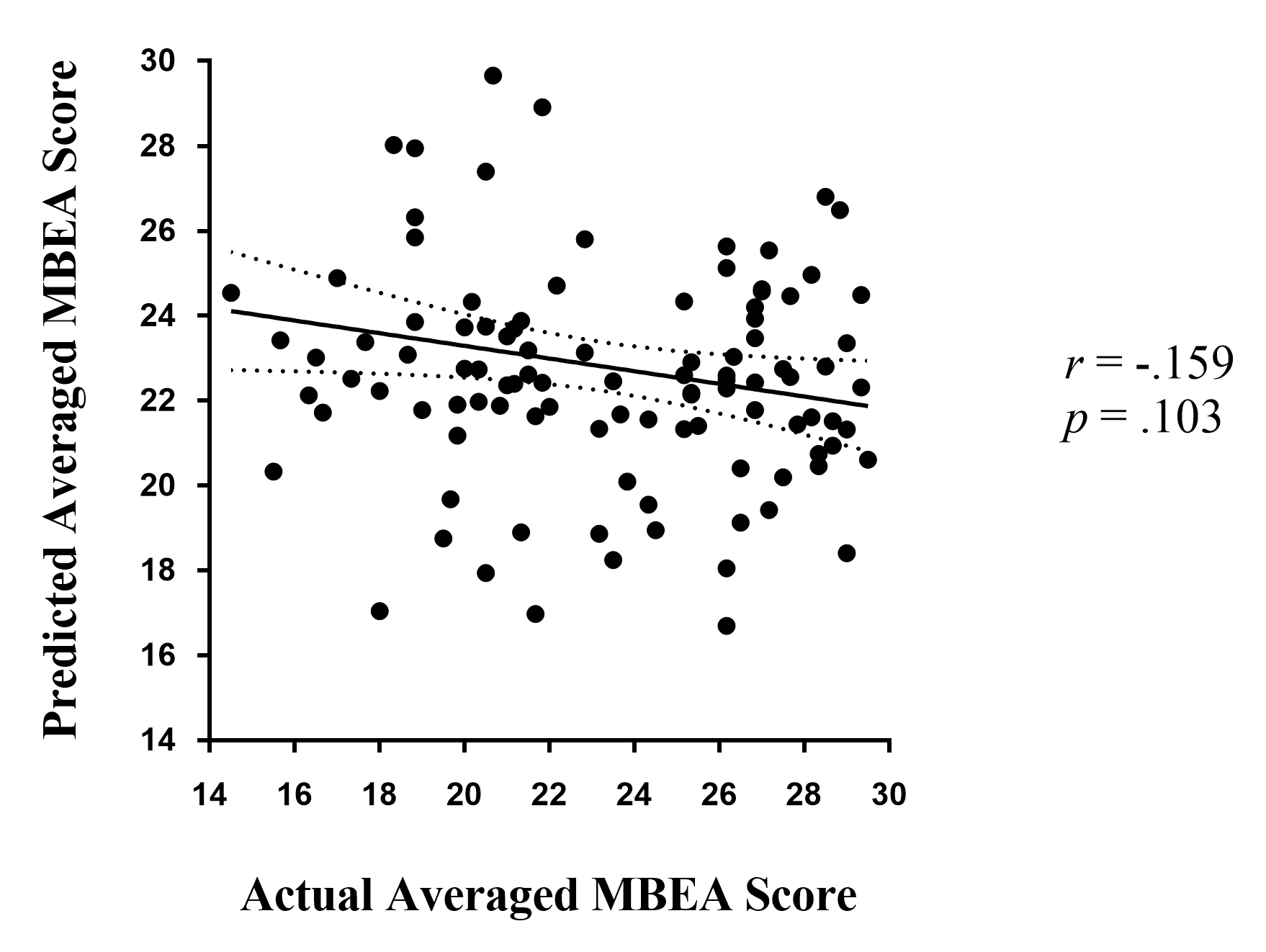

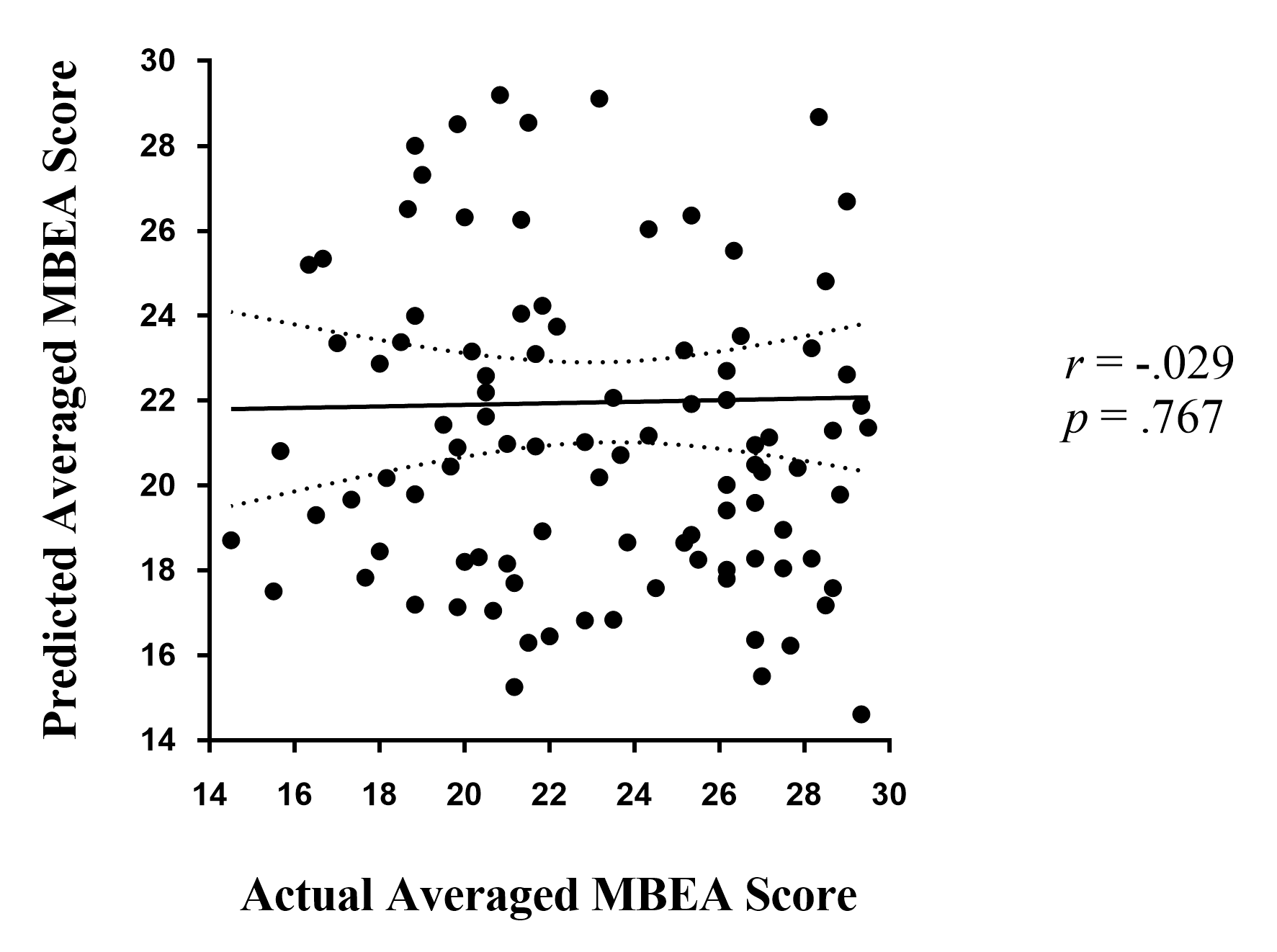

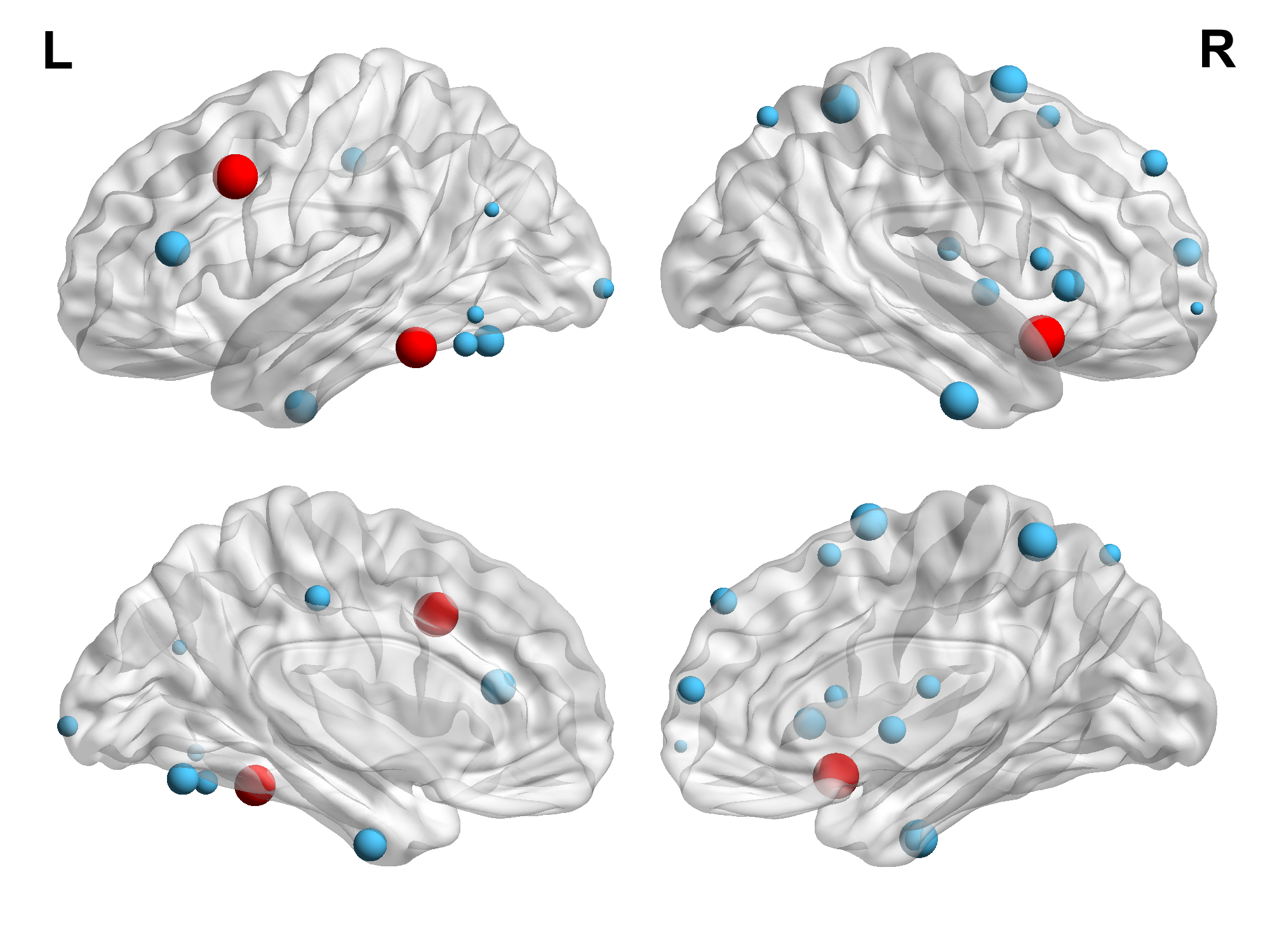

**Figure S2. All predictive ROIs when conducting prediction without covariates.** Greater volume represents greater contribution in prediction for each ROI. Most effective ROIs are marked with red color. Except R.ITG_A20il, most predictive ROIs are same with our main results.

Spearman Correlation: Structure-function Coupling

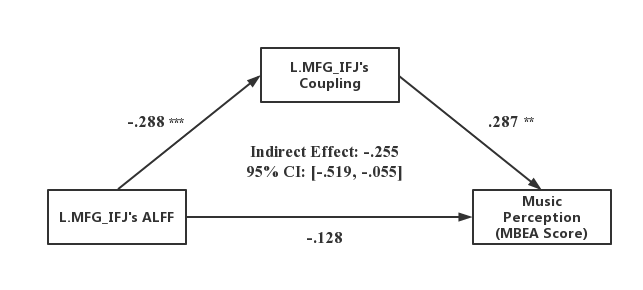
**Figure S3. Mediated relation among ALFF, structure-function coupling and MBEA score.** Structure-function coupling of L.MFG_IFJ fully mediated the relation between ALFF and MBEA score (partially standardized indirect effect = -.255).

**Table S1.** All predictive ROIs and their total weight, selected times and correlation coefficient across all 10 cross validations when covariates were ignored

| **ROI** | **Selected Times in CV** | **Total Weight** | **Mean Correlation R**  **(M ± SD)** |
| --- | --- | --- | --- |
| R.Ins_vIa | 10 | 115.139 | 0.254 ± 0.035 |
| L.MFG_IFJ | 10 | 106.076 | 0.277 ± 0.035 |
| L.ITG_A20cl | 10 | 61.708 | 0.250 ± 0.026 |
| R.SFG_A6dl | 7 | 36.196 | 0.229 ± 0.037 |
| R.ITG_A20il | 6 | 41.641 | 0.214 ± 0.031 |
| R.Tha_cTtha | 6 | 5.276 | -0.207 ± 0.034 |
| R.PCun_A5m | 5 | 48.045 | 0.199 ± 0.032 |
| L.IFG_IFS | 5 | 29.176 | 0.198 ± 0.035 |
| R.IFG_A44op | 3 | 19.671 | 0.172 ± 0.042 |
| L.FuG_A37mv | 3 | 17.411 | 0.194 ± 0.02 |
| R.SFG_A10m | 3 | 10.341 | -0.187 ± 0.034 |
| R.SFG_A9l | 3 | 8.646 | -0.176 ± 0.031 |
| L.PhG_A35/36r | 2 | 21.796 | 0.154 ± 0.044 |
| R.BG_dlPu | 2 | 9.619 | 0.177 ± 0.029 |
| L.CG_A23c | 2 | 7.306 | 0.187 ± 0.017 |
| R.SPL_A7c | 2 | 3.959 | 0.174 ± 0.032 |
| L.LOcC_OPC | 2 | 3.369 | -0.164 ± 0.032 |
| R.MFG_A10l | 2 | 0.569 | -0.166 ± 0.038 |
| L.ITG_A37elv | 1 | 6.132 | 0.115 ± 0.049 |
| R.SFG_A8m | 1 | 5.215 | 0.142 ± 0.042 |
| R.IFG_A44v | 1 | 5.106 | 0.138 ± 0.032 |
| L.ITG_A37vl | 1 | 1.775 | 0.141 ± 0.042 |
| L.IPL_A39rv | 1 | 1.077 | 0.136 ± 0.034 |

**Abbreviations:** ROI: Region of Interest; M ± SD: Mean ± Standard Deviation; CV: Cross Validation; Ins: Insula; MFG: Middle Frontal Gyrus; ITG: Inferior Temporal Gyrus; CG: Cingulate Gyrus; SFG: Superior Frontal Gyrus; IFG: Inferior Frontal Gyrus; Tha: Thalamus; FuG: Fusiform Gyrus; SPL: Superior Parietal Lobule; PCun: Precuneus; PhG: Parahippocampus Gyrus; BG: Basal Ganglia; LOcC: lateral Occipital Cortex; IPL: Inferior Parietal Lobule.

**Table S2.** Spatial correlation of R.ITG_A20il and cognitive terms in NeuroSynth

| R.ITG_A20il Only | | R.ITG_A20il with Connected Regions | |
| --- | --- | --- | --- |
| Cognitive Terms | Spatial Correlation | Cognitive Terms | Spatial Correlation |
| concentration | 0.061 | theory mind | 0.197 |
| prospective | 0.057 | social | 0.156 |
| expertise | 0.056 | mental states | 0.154 |
| intentions | 0.055 | theory | 0.141 |
| watching | 0.043 | face | 0.138 |
| theory mind | 0.030 | mind tom | 0.135 |
| social | 0.027 | mentalizing | 0.133 |
| light | 0.027 | face recognition | 0.129 |
| happy faces | 0.026 | social interaction | 0.119 |
| learn | 0.025 | visual | 0.115 |
| compulsive disorder | 0.024 | belief | 0.109 |
| experiences | 0.022 | recognize | 0.109 |
| atrophy | 0.021 | social cognitive | 0.108 |
| videos | 0.021 | motion | 0.108 |
| obsessive | 0.020 | extrastriate | 0.103 |
| amnestic | 0.020 | beliefs | 0.101 |
| obsessive compulsive | 0.020 | psts | 0.100 |
| personality | 0.018 | intentions | 0.100 |
| instructions | 0.018 | viewing | 0.100 |
| mild cognitive | 0.017 | theory mind | 0.197 |
| major depression | 0.016 | social | 0.156 |
| recall | 0.016 | mental states | 0.154 |
| parkinson disease | 0.015 |  |  |

**Table S3.** Spatial correlation of R.Ins_vIa and cognitive terms in NeuroSynth

| R.Ins_vIa Only | | R.Ins_vIa with Connected Regions | |
| --- | --- | --- | --- |
| Cognitive Terms | Spatial Correlation | Cognitive Terms | Spatial Correlation |
| inhibitory control | 0.169 | pain | 0.081 |
| inhibition | 0.166 | listening | 0.078 |
| response inhibition | 0.106 | stop | 0.074 |
| taste | 0.071 | mood | 0.074 |
| emotional information | 0.070 | reward | 0.072 |
| stop | 0.059 | gain | 0.065 |
| error | 0.054 | response inhibition | 0.064 |
| stop signal | 0.051 | motivation | 0.064 |
| noxious | 0.041 | intensity | 0.063 |
| anxiety | 0.039 | inhibition | 0.062 |
| reactivity | 0.037 | music | 0.060 |
| reward | 0.037 | taste | 0.058 |
| disgust | 0.036 | disgust | 0.057 |
| risk taking | 0.036 | anticipation | 0.056 |
| emotional | 0.032 | motivational | 0.054 |
| fear | 0.032 |  |  |
| losses | 0.030 |  |  |
| threatening | 0.030 |  |  |
| heart | 0.029 |  |  |
| autonomic | 0.029 |  |  |
| gain | 0.028 |  |  |
| painful | 0.025 |  |  |
